# Spatial Attention Enhances the Neural Representation of Invisible Signals Embedded in Noise

**DOI:** 10.1101/102731

**Authors:** Cooper A. Smout, Jason B. Mattingley

## Abstract

Recent evidence suggests that voluntary spatial attention can affect neural processing of visual stimuli that do not enter conscious awareness (i.e. invisible stimuli), supporting the notion that attention and awareness are dissociable processes (Watanabe et al., 2011; Wyart, Dehaene, & Tallon-Baudry, 2012). To date, however, no study has demonstrated that these effects reflect enhancement of the neural representation of invisible stimuli *per se*, as opposed to other neural processes not specifically tied to the stimulus in question. In addition, it remains unclear whether spatial attention can modulate neural representations of invisible stimuli in direct competition with highly salient and visible stimuli. Here we developed a novel electroencephalography (EEG) frequency-tagging paradigm to obtain a continuous readout of human brain activity associated with visible and invisible signals embedded in dynamic noise. Participants (*N* = 23) detected occasional contrast changes in one of two flickering image streams on either side of fixation. Each image stream contained a visible or invisible signal embedded in every second noise image, the visibility of which was titrated and checked using a two-interval forced-choice detection task. Steady-state visual-evoked potentials (SSVEPs) were computed from EEG data at the signal and noise frequencies of interest. Cluster-based permutation analyses revealed significant neural responses to both visible and invisible signals across posterior scalp electrodes. Control analyses revealed that these responses did not reflect a subharmonic response to noise stimuli. In line with previous findings, spatial attention increased the neural representation of visible signals. Crucially, spatial attention also increased the neural representation of invisible signals. As such, the present results replicate and extend previous studies by demonstrating that attention can modulate the neural representation of invisible signals that are in direct competition with highly salient masking stimuli.

## Introduction

When viewing a cluttered visual scene, representations of the various objects compete for limited neural resources (Broadbent, 1958; Desimone & Duncan, 1995). Such ongoing neural competition can be biased by top-down mechanisms to facilitate the observer’s behavioural goals (Beck & Kastner, 2009). For example, voluntarily allocating covert spatial attention to a specific region of the visual field can selectively boost neural representations of salient stimuli within that region (Hillyard & Anllo-Vento, 1998; Martinez et al., 1999; Müller et al., 1998). Interestingly, recent studies demonstrate that spatial attention can also affect neural processing of weak stimuli that do not enter awareness (equated here with the contents of conscious experience; Schurger et al., 2008; Wyart and Tallon-Baudry, 2008; Wyart et al., 2012). However, since attention encompasses a variety of neural mechanisms (for a review see Womelsdorf and Everling, 2015), it remains unclear which subcomponents activate during processing of invisible stimuli. In particular, no study to date has tied neural activity to specific invisible stimuli, and thus it remains unclear whether spatial attention enhances *neural representations* of invisible stimuli or merely activates other neural mechanisms not specific to neural representations *per se* (e.g. alerting, orienting, or suppression mechanisms). Evidence that spatial attention increases the neural representation of invisible stimuli, without a corresponding increase in object awareness, would provide clear evidence that attention and awareness dissociate at the level of stimulus representations. Furthermore, previous studies presented invisible stimuli at different times or locations to highly visible masking stimuli, and thus it remains unclear how spatial attention treats neural representations of invisible signals that are in direct competition with visible stimuli. Such research is necessary if we are to understand how top-down mechanisms in the visual system allocate limited resources to competing stimuli with different levels of bottom-up signal strength (i.e. salience). In the present study, we used electroencephalography (EEG) to measure neural representations of visible and invisible stimuli embedded in highly salient noise, and assessed the effect of voluntary covert spatial attention on these neural representations.

To investigate these questions, it is necessary to disambiguate relatively weak neural activity arising from subjectively invisible targets from the stronger responses associated with highly salient and spatially coincident masking stimuli. To date, however, no such technique has been devised to effectively distinguish the neural signatures of these weak and strong sensory inputs. If a train of stimuli is presented at a fixed frequency, however, a stable oscillatory response is produced in the brain that can be observed in the frequency-domain in EEG recordings (the steady-state visual-evoked potential; SSVEP; Regan, 1966). Multiple stimuli in a visual scene can thus be ‘frequency tagged’ when flickered at unique frequencies, an approach that has proven useful for exploring the effects of attention on visible stimuli at separate spatial locations (Norcia, Appelbaum, Ales, Cottereau, & Rossion, 2015).

A recent study by Ales et al. (2012) pioneered a novel SSVEP technique for measuring neural representations of signals embedded in dynamic noise. In their study, Ales et al. presented participants with streams of luminance- and amplitude-matched noise images at a rate of 6 Hz. Every second image contained a face stimulus embedded in noise, and the coherence of the face was gradually increased over the duration of the trial until participants indicated they had detected it. Crucially, power at the frequency of signal presentation (3 Hz, representing the face in every second image) was found only in trials that contained embedded faces, and not in trials in which the face was replaced by another noise display. Thus, the neural activity at the frequency of the embedded signal serves as a useful measure of the neural representation of that stimulus, irrespective of any other neural processes that may be operating concurrently.

Using the same principle as Ales et al. (2012), we developed a novel paradigm to obtain a continuous readout of neural activity associated with visible and invisible signals embedded in dynamic noise. Participants directed attention to one of a pair of flickering image streams to detect occasional contrast changes, and we assessed the effect of spatial attention on neural representations of both visible and invisible signals. We employed a two-interval, forced-choice signal detection task to confirm that appropriate levels of signal coherence were selected for visible and invisible signals. To anticipate, we found that spatial attention enhanced neural representations of both visible and invisible signals, suggesting that attention can bias neural activity in favour of invisible stimuli that are in spatial and temporal competition with highly salient masking noise.

## Materials and Methods

### Participants

Twenty-three healthy participants (11 female, mean age: 22.65 years) with normal or corrected-to-normal vision were recruited via an online research participation scheme at The University of Queensland. Participants completed a safety-screening questionnaire and provided written informed consent prior to commencement of the study, which was approved by The University of Queensland Human Research Ethics Committee.

### Stimuli and apparatus

The method of stimulus generation (Figure 1) was adapted from Ales, Farzin, Rossion and Norcia (2012) to maintain the same average power distribution and luminance across all images. All images were created from the same seed image consisting of an annulus (seven cycles, inner diameter: 4.67° of visual angle, outer diameter: 14° of visual angle) on a uniform mid-grey square background (14° of visual angle; Figure 1a, top left). The phase distribution of the seed image was randomised to create a noise background with the same amplitude distribution as the seed image (Figure 1a, bottom left). The annulus and noise background were then combined using complementary spatial blending masks (which spanned from the annulus edges to 2° of visual angle within each edge; Figure 1a, top and bottom right) to create an exemplar image consisting of a fully coherent annulus on a noise background (Figure 1a, center right). The phase distribution of this exemplar image was then ‘scrambled’ (randomized) to the extent required by the trial sequence (see *Stimulation Protocol,* below): phase angles of ‘noise images’ were scrambled completely (Figure 1b, bottom), whereas phase angles of ‘signal images’ were linearly interpolated between the original phase angles and a random phase distribution (Figure 1b, top and middle). Because phase angles are circular, interpolation of phase angles was computed in the direction of least difference to maintain a uniform phase distribution (Ales, Farzin, & Norcia, 2012).

**Figure 1.**
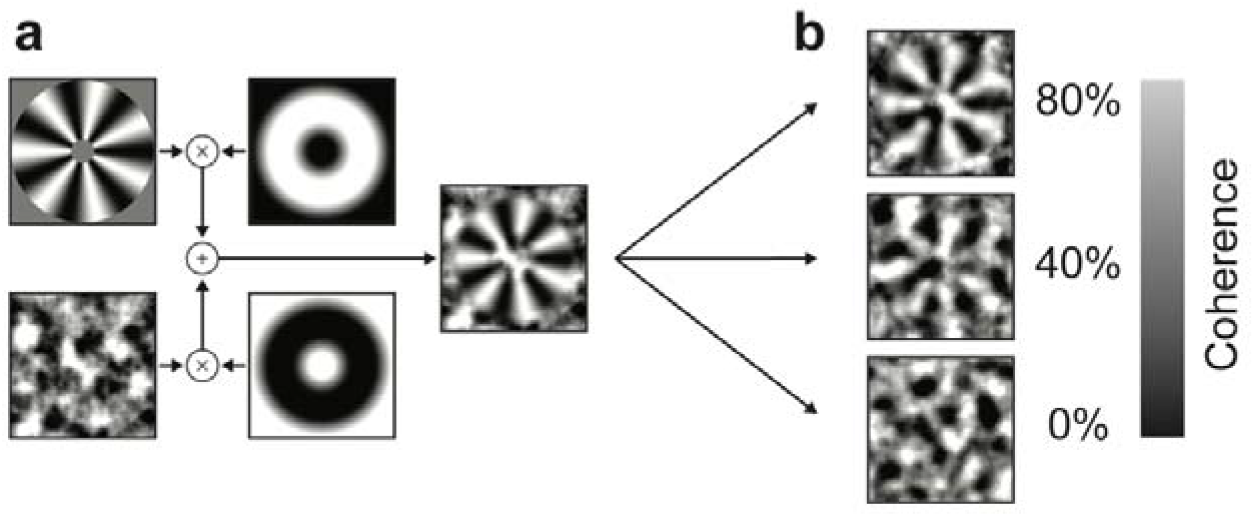
Stimulus generation. **(*a*)** Phase distribution of the signal (annulus, top left) was scrambled to create a noise background that was different for every image (bottom left). Signal and noise images were combined via inverse masks to create an exemplar image (centre right), which was then phase-scrambled according to the desired level of signal coherence. **(*b*)** Example images containing visible signal (top), invisible signal (middle), or noise only (bottom).

Thus, all images contained some amount of ‘noise’, which represented the (partially or completely) randomized phase distribution of its exemplar image. Signal images contained noise both ‘behind’ the annulus (in the exemplar background), as well as ‘in front of’ the annulus, since the phase-structure of the exemplar image was partially randomized in the final image creation step. Since each exemplar image was created using a unique noise background, the only consistent structure between any two images was the signal itself, subject to its level of phase coherence. Furthermore, since all images – both signal and noise - were created from the same seed annulus image, all images in the experiment shared the same low-level characteristics, including amplitude and luminance.

Stimuli were presented on a 21-inch CRT monitor (NEC, Accusync 120) with a screen resolution of 800 × 600 pixels and a refresh rate of 120 Hz, using the Cogent 2000 Toolbox (http://www.vislab.ucl.ac.uk/cogent.php) for Matlab (The Mathworks Inc., Natick, USA) running under Windows XP. Participants were seated in a comfortable armchair in an electrically shielded laboratory, with the head supported by a chin rest at a viewing distance of 57cm.

### Procedures

The present study used a within-participant design with two levels of target awareness (*visible, invisible*) and two levels of spatial attention (*attended, ignored).* Two tasks with similar overall designs were employed to manipulate awareness and spatial attention.

#### Awareness Task

Participants were presented with two flickering image streams on either side of fixation (visual angle: 14°), as shown in Figure 2 and *Movie 1.* Each image stream contained two consecutive intervals of 2.4 *s* duration (see *Stimulation Protocol* for interval details). One of the intervals in each image stream (randomized separately) contained signal (the ‘signal interval’) and the other interval contained noise only (the ‘noise interval’). Participants were asked to maintain fixation and report, on the cued side, which of the two intervals contained signal (two-interval forced-choice), while ignoring the non-cued side. The cue direction (left or right) was randomized for the first trial of each block and then alternated every eight trials.

**Figure 2.**
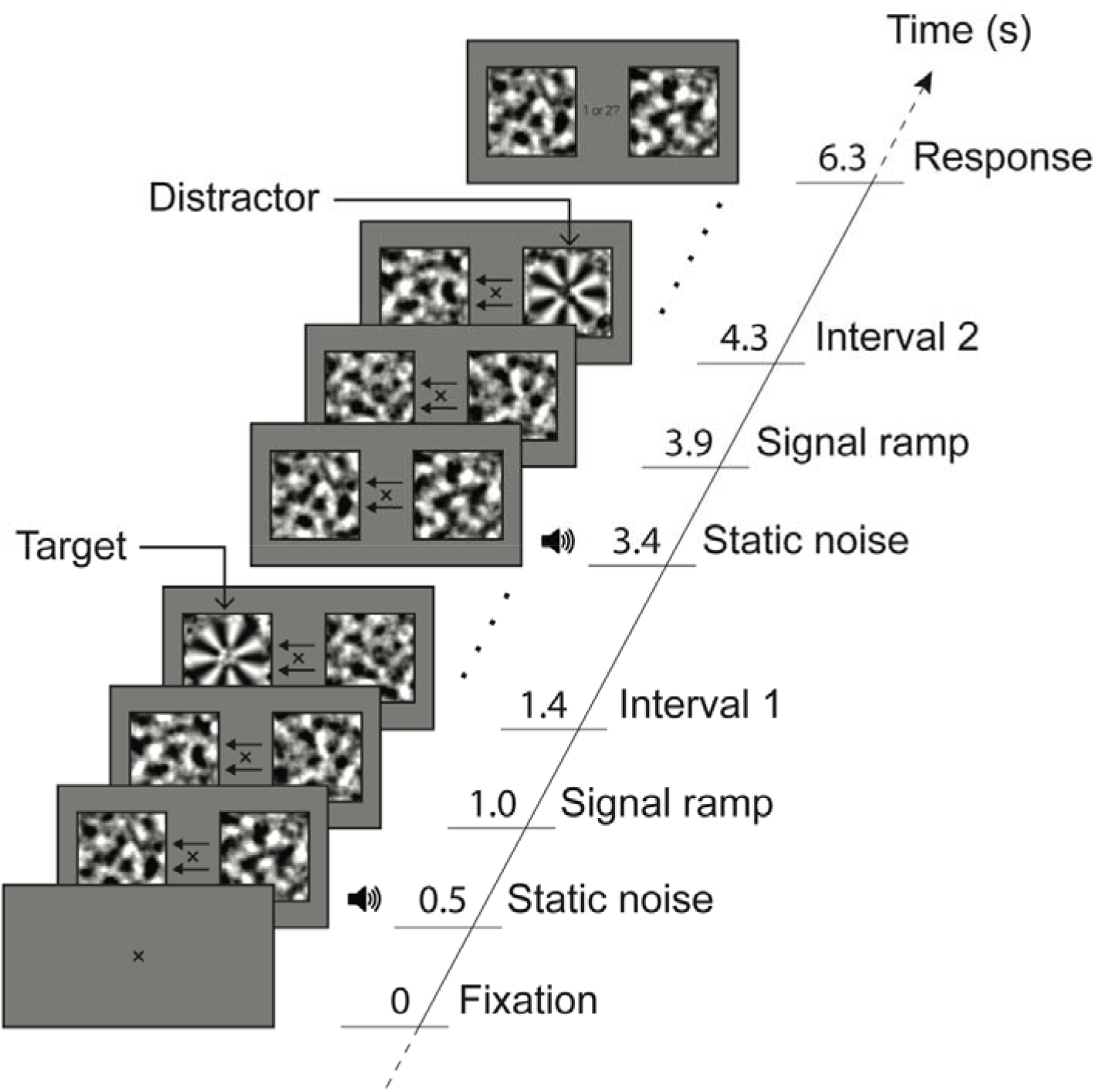
Awareness Task. Participants fixated centrally and searched for a signal embedded in dynamic noise on the cued side, which appeared in only one of two consecutive intervals. In the example shown, a target is present during interval 1 on the cued (left) side. Note that a distractor signal is also present during interval 2 on the ignored (right) side. Images flickered during the ramping and signal intervals only (see Figure 1b for typical image sequence).

Participants completed two versions of the Awareness Task. The first version was run at the beginning of the experiment (following practice with accuracy feedback), in order to set signal coherence levels for the subsequent Attention Task (see below). In this first version, participants completed 48 trials with feedback, while levels of signal coherence were adjusted according to an adaptive Quest staircase (Watson & Pelli, 1983) designed to approximate the maximum level of signal coherence that could not be detected by each participant (i.e. the invisible condition). Signal coherence for the visible condition was then set 40% higher than this level, as guided by psychometric functions fitted to pilot data. The second version of the Awareness Task was run at the end of the experiment, to verify that appropriate levels of signal coherence had been selected. In this version, participants completed two blocks of 64 trials (without feedback), with each image stream containing visible *or* invisible signal in one of the two consecutive intervals (randomized separately across trials).

#### Attention Task

Participants were again presented with two flickering image streams on either side of fixation, as shown in Figure 3 and *Movie 2.* Unlike in the Awareness Task, however, in the Attention Task each image stream contained only one interval of 10 *s* duration per trial, and both image streams contained either visible or invisible signals (as per the staircase procedure above). Additionally, each image stream occasionally decreased in contrast before returning to normal across a 1 *s* period (ramping on and off linearly), with at least 1.5 *s* between peaks of contrast decreases (in either stream). Participants were asked to maintain fixation and report at the end of the trial how many contrast decreases (*targets*) occurred in the cued (*attended*) image stream. When the attended stream contained two contrast targets, the second target peaked between 7 *s* and 8.5 *s* into the trial, to encourage sustained attention throughout trials. Participants were allowed to practice the task (with feedback after each trial) before completing eight blocks of 64 test trials, with feedback provided between blocks. The percentage of contrast decrease was adjusted between blocks to maintain an approximate detection level of 65% (according to a 1 up / 2 down staircase with step sizes of 5%).

**Figure 3.**
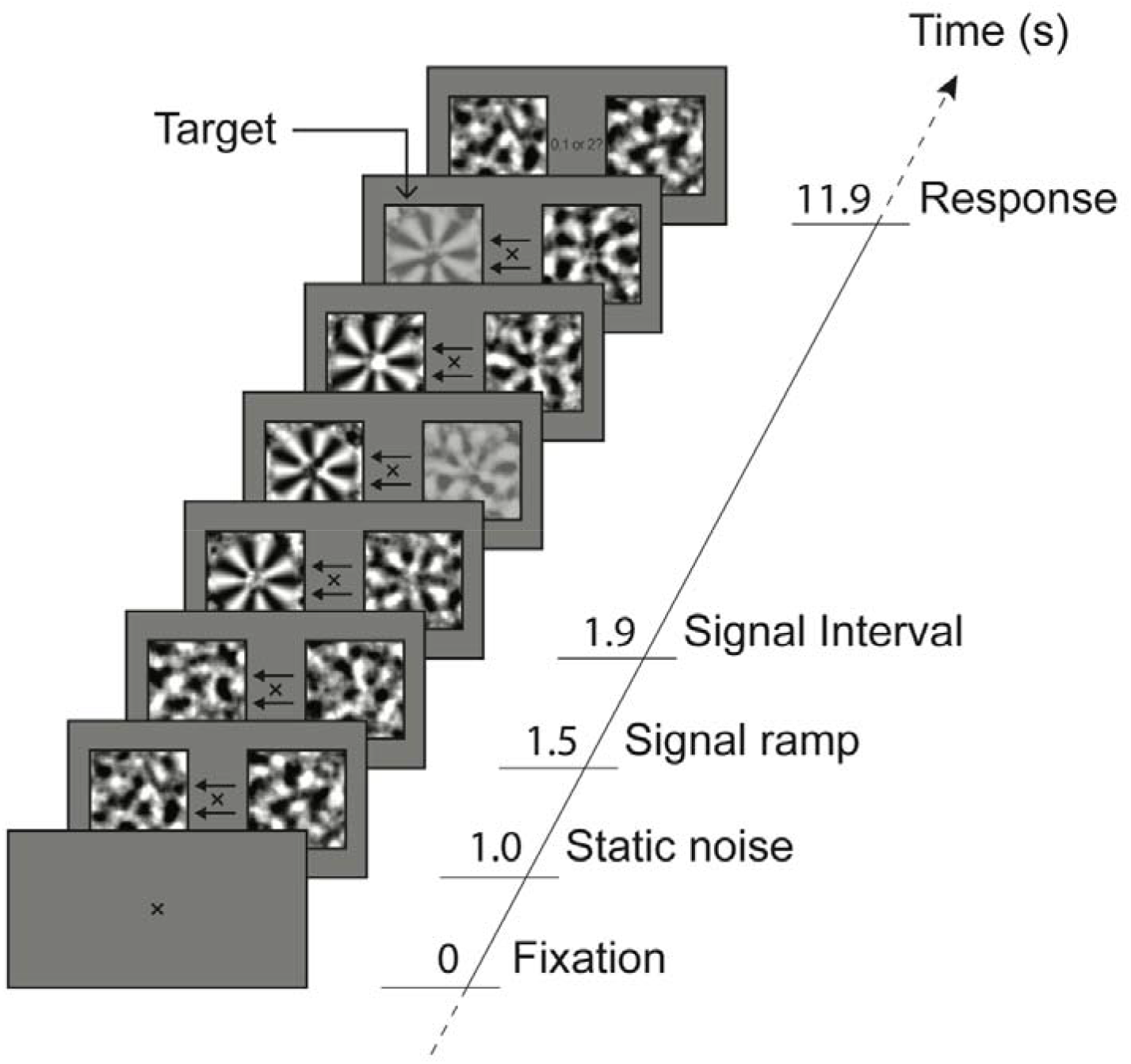
Attention Task. Participants fixated centrally and counted the number (0, 1 or 2) of brief decreases in contrast in the cued (attended) image stream. In the example shown, one contrast decrease appeared in each of the attended (left) and ignored (right) image streams. Each image stream contained a visible or invisible annulus embedded in dynamic noise throughout the entire signal interval. Note that for illustrative purposes the magnitude of the contrast decrements has been enhanced in the figure.

#### Stimulation Protocol

During any one trial, intervals in the left and right image streams flickered at unique frequencies (10 and 15 Hz, counterbalanced across trials). Although Awareness Task trials contained two intervals per image stream and Attention Task trials contained only one interval per image stream, the structure of intervals in both tasks was essentially the same. Figure 4 shows the stimulation protocol for one interval flickering at 10 Hz. All intervals began with 0.5 s of static noise, after which images flickered consecutively at the designated frequency (10 or 15 Hz). The phase distributions of all images in ‘noise intervals’ (Awareness Task) were completely scrambled (see Stimuli and apparatus). During ‘signal intervals’ (Awareness and Attention Tasks), images alternated between completely phase-scrambled images (noise) and partially phase-scrambled images (signal). The coherence of signal images ramped up linearly during the first 0.4 *s* of signal intervals to eliminate involuntary capture of attention (Figure 4). At the end of the flicker duration (2.4 *s* for Awareness Task trials, 10.4 *s* for Attention Task trials), static noise was presented until the next interval began flickering (first interval of Awareness Task trials only) or the participant responded (Attention Task trials and second interval of Awareness Task trials).

**Figure 4.**
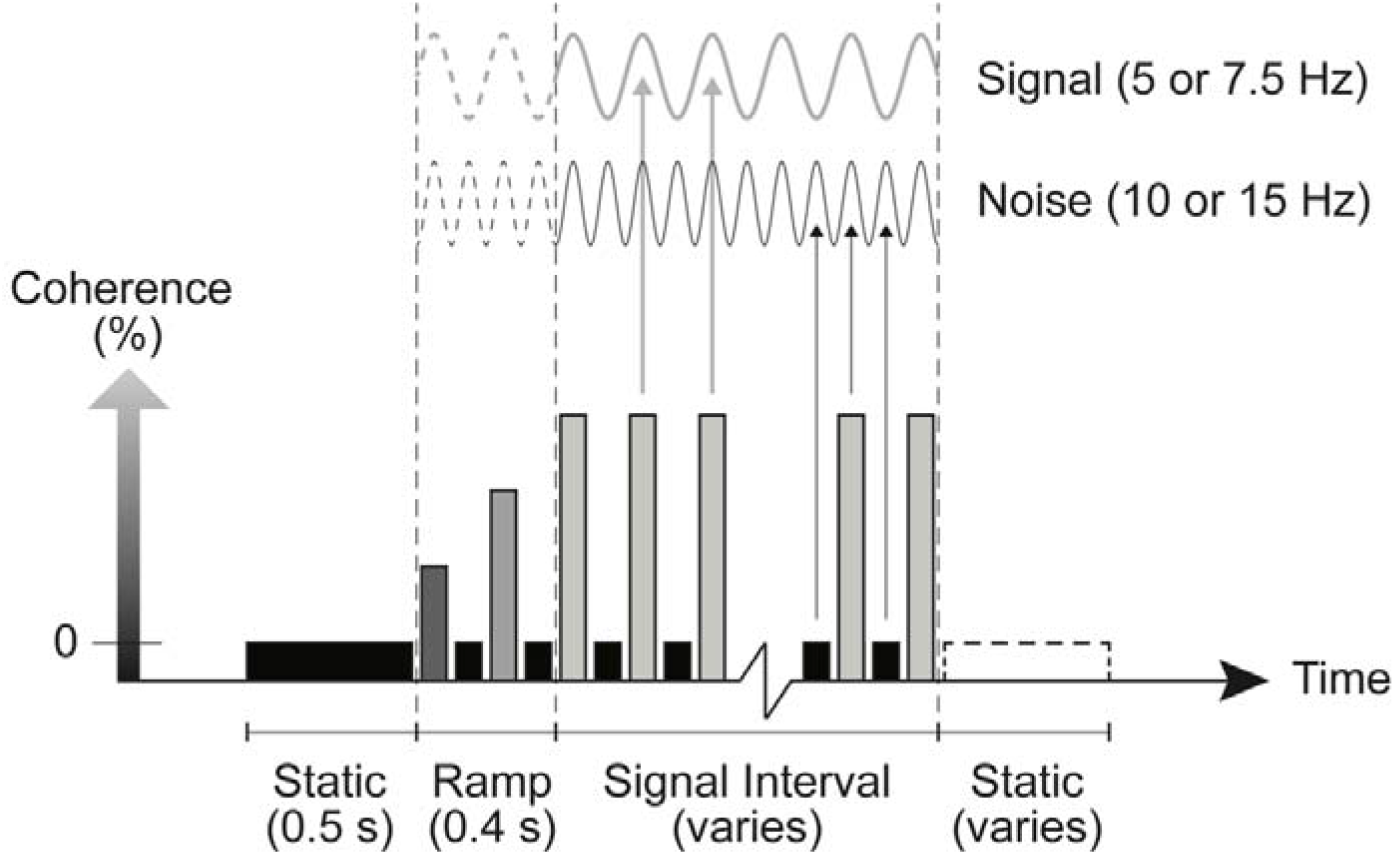
Schematic showing dynamic change in signal coherence during one interval of a single trial in which displays flickered at 10 Hz. Black bars represent images that were completely phase-scrambled (i.e., noise images with zero signal coherence), and light grey bars represent images that retained some level of signal structure (coherence) after the phase-scrambling procedure (Figure 1b). The coherence of signal images increased linearly during the ramping period (i.e., the first 0.4 *s* of each flickering interval), then remained at the specified level until the interval end (i.e., after 10 *s* in the Attention Task, and 2.4 *s* in the Awareness Task). Shown above the image sequence are putative neural responses driven by the signal and noise stimuli at distinct frequencies. Note that all images contain some amount of noise, and therefore contribute to the neural response at noise frequencies (10 Hz in this example, though in 50% of trials the noise frequency was 15 Hz). Only every second image contained consistent structure (the signal), and these images therefore contributed to neural responses at half the noise frequency (5 Hz in this example, but 7.5 Hz in the other 50% of trials).

Shown at the top of Figure 4 are putative neural responses evoked by the stimulation protocol. Since all flickering images contained some amount of ‘noise’, SSVEP responses were expected to be elicited by noise stimuli at the *noise frequency* (i.e., 10 or 15 Hz). Crucially, since a signal was embedded in every second image during signal intervals, a separate SSVEP was expected to be elicited at half the noise frequency in response to signal (5 or 7.5 Hz, the *signal frequency).* Thus, we were able to isolate neural responses to both noise and signal (at two levels of awareness) when those stimuli were either attended or ignored (see Results for details of power computation).

### EEG recording

Participants were fitted with a 64 Ag-AgCl electrode EEG system (BioSemi Active Two: Amsterdam, Netherlands) after the initial Awareness Task, and EEG data were recorded during the Attention Task and final Awareness Task. Continuous data were recorded using BioSemi ActiView software (http://www.biosemi.com), and were digitized at a sample rate of 1024 Hz with 24-bit A/D conversion and a .01 – 208 Hz amplifier band pass. All scalp channels were referenced to the standard BioSemi reference electrodes, and electrode offsets were adjusted to be below 25 μV before beginning the recording. Horizontal and vertical eye movements were recorded via pairs of BioSemi flat Ag-AgCl electro-oculographic electrodes placed to the outside of each eye, and above and below the left eye, and respectively.

### EEG data pre-processing

Electroencephalography (EEG) recordings were processed offline using the Fieldtrip toolbox in Matlab (http://fieldtrip.fcdonders.nl). Trials containing horizontal eye movements were inspected manually and rejected if lateral eye fixations exceeded 1 *s* during the Attention Task (3.55% of trials) or 150ms during the final Awareness Task (12% of trials). Two faulty electrodes (across two participants) were interpolated using the nearest neighboring electrodes. Scalp electrode data were re-referenced to the average of all 64 electrodes, resampled to 256 Hz, and subjected to a surface Laplacian filter (M. Cohen, 2014). Trials were epoched into intervals containing signal at full coherence (Awareness Task: 1.4 – 3.4 *s* or 4.3 – 6.3 *s*, Figure 2; Attention Task: 1.9 *s* – 11.9 *s*, Figure 3), for frequency power analyses (see Results). Attention Task trials were also epoched with an additional 2 *s* before and after each signal period for time-frequency power analyses.

### Phase-locked Power Calculation

To measure neural responses to flickering stimuli in the Attention and Awareness Tasks, we examined *phase-locked power* (sometimes called ‘evoked’ power) at each of the noise (10 and 15 Hz) and signal (5 and 7.5 Hz) stimulation frequencies. We elected to use phase-locked power as our measure of interest because it is maximally sensitive to neural responses in phase with the events of interest - in our case the onsets of flickering images - and parcels out these responses from non-phase-locked neural activity (sometimes called ‘induced’ power) that might otherwise obscure weak neural responses to invisible signals.

Phase-locked power was calculated as the difference between normalized *total power* and *non-phase-locked power* (Cohen, 2014). Total raw power was computed by applying Fourier transforms (Hanning window, 0.10 Hz frequency resolution) to 10 *s* trial epochs in the Attention Task (1.9 - 11.9 *s*: Figure 3) and 2 *s* interval epochs in the Awareness Task (1.4 – 3.4 *s* and 4.3 – 6.3 *s*, Figure 2; zero-padded to 10 s), and averaging across trials in each condition of interest (attention, awareness, stimulation frequency and side). Total power in each condition was then decibel-normalized by dividing the raw power in each frequency bin by the average power in the 20 adjacent frequency bins (+/- 1.0 Hz) and multiplying the logarithmic transform of the result by 10 (M. Cohen, 2014). Non-phase-locked power was calculated in the same manner as total power, after the condition-average event-related potential had been subtracted from each trial (M. Cohen, 2014). Finally, phase-locked power (hereafter referred to as *power*) was calculated by subtracting the non-phase-locked power from the total power within each condition.

To test whether participants maintained covert attention during the Attention Task, we also calculated noise frequency power as a function of time. Preprocessed EEG data were bandpass filtered at each frequency of interest (Matlab function: fir1, order: 64 samples, width: .01 Hz), subjected to a Hilbert transform, and down-sampled to 40 Hz. Phase-locked time-frequency power was then calculated in the same manner as phase-locked frequency power (above).

To maximise power for all statistical analyses, we subjected the data to a contralateralization procedure to remove the side of stimulation (left or right of fixation) as a factor within each attention and awareness condition. The electrode labels in trials with right-sided stimulation (i.e., when stimuli on the right of fixation flickered at the frequency of interest) were mirrored along the sagittal centre-line (e.g., PO7 became PO8, and vice versa). After this procedure, left-sided electrodes in all trials (irrespective of stimulation side) represented those ipsilateral to stimulation, and right-sided electrodes represented those contralateral to stimulation. Since hemispheric differences were not crucial to our research question, we then collapsed power across the factor of stimulation side. All electrode topographies presented here (Figures 6, 7, 8, and 10) represent data that underwent this contralateralization procedure.

**Figure 6.**
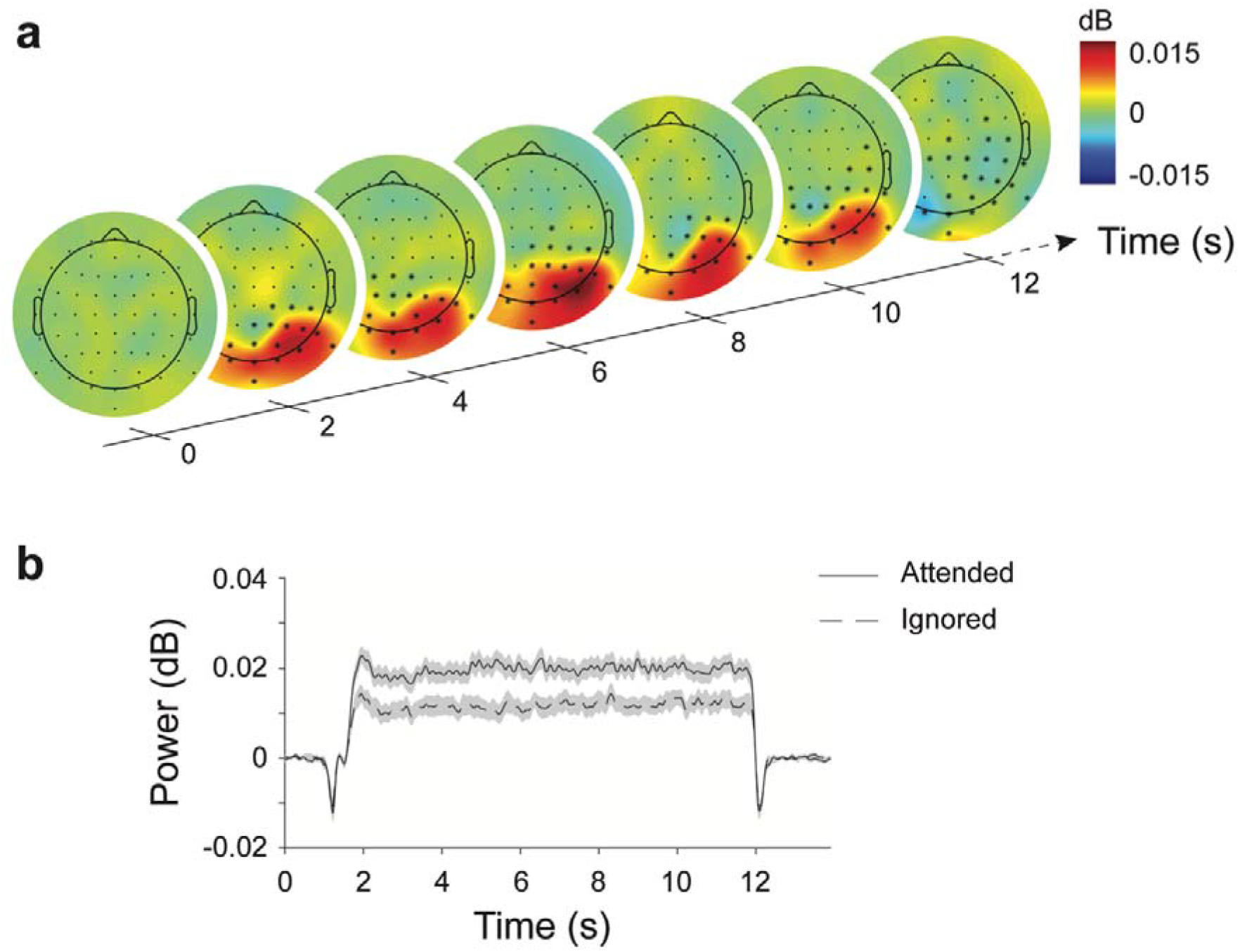
Effect of spatial attention on the neural response to noise in the Attention Task. **(*a*)** Electrode topographies represent the difference between attended and ignored noise SSVEPs, contralateralized to represent left side stimulation, and collapsed across noise frequencies (10 and 15 Hz). Stars indicate the cluster of electrodes that showed significantly greater noise frequency power with attention within +/− 1 *s* of the displayed timepoint (cluster-corrected *p* < .001). **(*b*)** Phase-locked power averaged across contralateral electrodes P1/2, P3/4, P5/6, P7/8, P9/10, PO3/4, PO7/8, and O1/2. Shaded regions indicate the within-subjects standard error of the mean.

## Results

### Awareness Task

The initial adaptive staircase procedure produced an average signal coherence of 29.91% (SD = 3.18%) for the invisible condition and 69.91% (SD = 3.18%) for the visible condition, across participants. One-tailed t-tests were used to assess signal awareness in the final Awareness Task, which revealed that visible targets were detected above chance (chance = 50%; M = 95.77%, SD = 3.64%, *t*_(22)_ = 60.367, *p* < .001) and that invisible targets were detected no better than chance (M = 50.96%, SD = 8.13%, *t*_(22)_ = .565, *p* = .289). Furthermore, Bayesian statistics supported the null hypothesis that invisible stimuli were detected at chance (uniform prior, lower bound = 50%, upper bound = 100%, *B* = .07).

### Attention Task

One-tailed t-tests revealed that contrast decrement targets were detected better than chance level (chance = 33%; M = 65.72%, SD = 6.77%, *t*_(22)_ = 46.302, *p* < .001). A two-tailed t-test revealed that contrast decrement targets were better detected when the signal was visible (M = 68.11%, SD = 8.38%) than when it was invisible (M = 63.34%, SD = 5.71%, *t*_(22)_ = 4.84, *p* < .001).

### Noise and Signal Elicit Distinct Neural Respomes

To confirm that our measure of phase-locked power successfully isolated neural responses to signal and noise stimuli, we computed power in the Attention Task (see Methods) and collapsed across awareness conditions and participants. Figure 5 shows the phase-locked power at contralateral electrode PO3/4 as a function of frequency, separately for each combination of stimulation frequencies. Note that power is only greater than zero at the signal (5 and 7.5 Hz) and noise (10 and 15 Hz) frequencies, confirming that the measure successfully isolated neural responses to the flickering stimuli.

**Figure 5.**
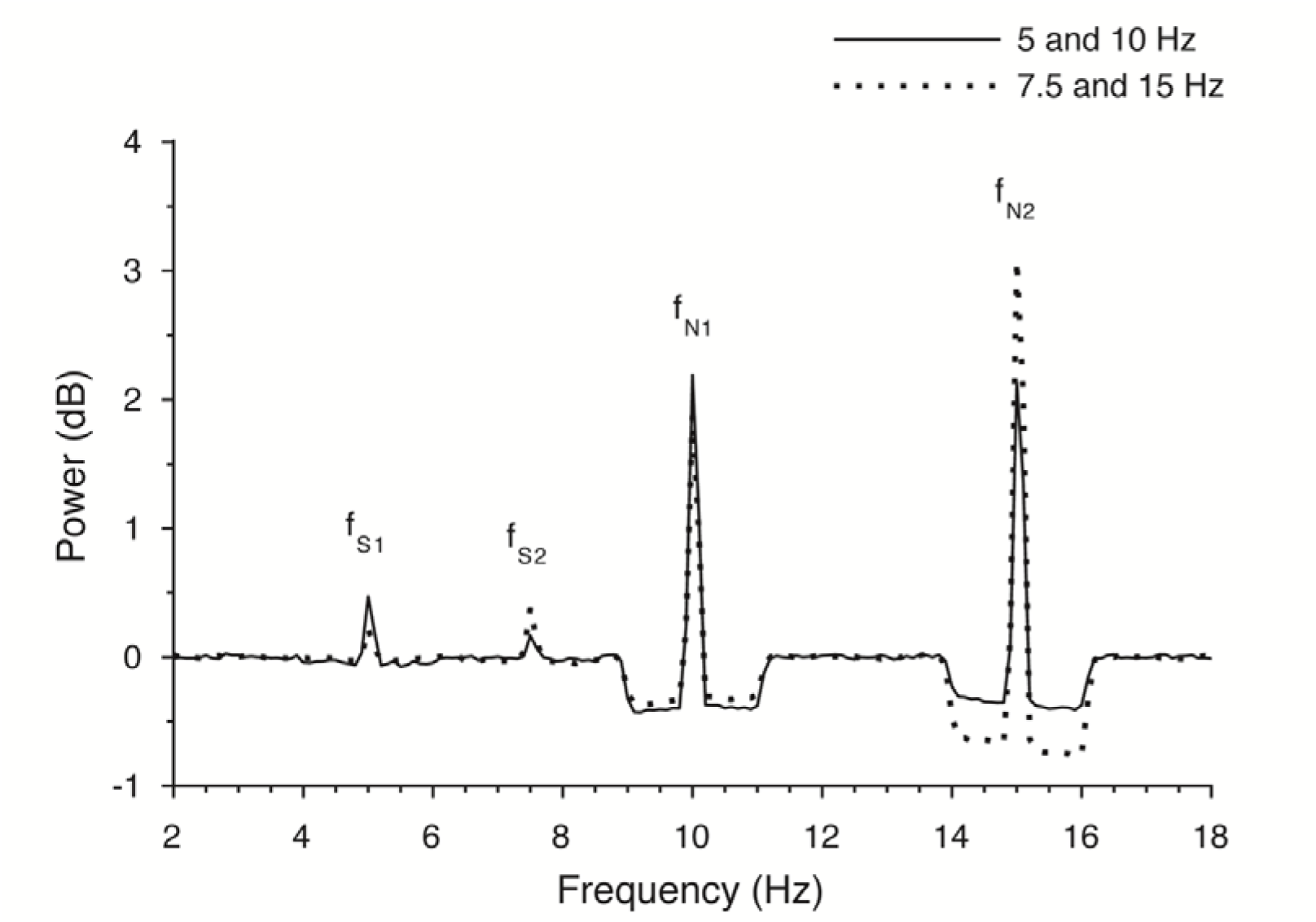
Phase-locked power at contralateral electrode PO3/4 in the Attention Task, averaged across all conditions and participants. Note that peaks in the frequency spectrum only occur at the signal (5 and 7.5 Hz) and noise frequencies (10 and 15 Hz).

### Spatial Attention Enhances Neural Representations of Noise

To verify that covert attention was directed to the cued image stream (left or right) throughout Attention Task epochs, we assessed differences in time-frequency power between attended (cued) and ignored image streams. Time-frequency power was computed using Hilbert transforms (see Methods) and collapsed across noise frequencies and awareness conditions (since all stimuli contained noise). The effect of attention was then tested with a two-tailed Monte-Carlo cluster-based permutation test in the *Fieldtrip* toolbox for Matlab (between participant factors: electrode power and time, cluster p < .05, unit p < .05, 1000 permutations). Cluster-based permutation analyses are a non-parametric method for testing condition differences in high-dimensional neural data, while correcting for multiple comparisons (for a detailed discussion, see Maris & Oostenveld, 2007). They are typically most useful when experimenters have few *a priori* expectations about specific locations or times of effects (Groppe, Urbach, & Kutas, 2011), as was the case in the current investigation. As revealed in Figure 6, spatial attention enhanced noise frequency power across a cluster of posterior and contralateral electrodes that spanned the entire epoch (cluster-corrected *p* = .002).

### Target Detection Correlates with the Effect of Attention on Neural Representations of Noise

Next, we investigated the relationship between behavioural performance on the Attention Task and the effect of attention on neural representations of noise stimuli. We labelled trials in which participants identified the exact number of targets (0, 1 or 2) as correct and all other trials as incorrect, and then balanced the number of correct and incorrect trials in each condition by removing a random subset of trials in the category with the greater number of trials. Noise frequency power was computed (see Methods) and collapsed across frequencies and sides, and the effect of attention was computed as the difference between attended and ignored trials (attended – ignored). Finally, the attentional modulation of correct and incorrect trials was compared with a two-tailed Monte-Carlo cluster-based permutation test (between participant factor: electrode power, cluster *p* < .05, unit *p* < .05, 1000 permutations). As can be seen in Figure 7, there was a larger effect of attention on the neural response to noise stimuli across frontal and central electrodes when targets (contrast decrements) were correctly detected (cluster-corrected *p* = .014).

**Figure 7.**
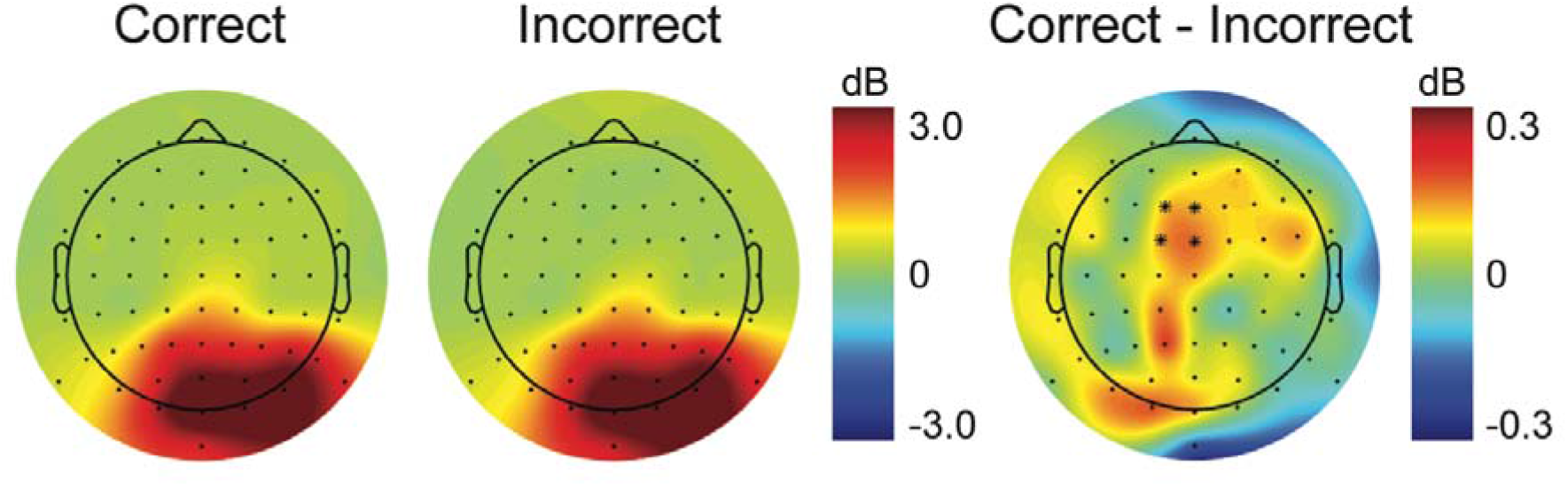
Relationship between target detection and the effect of attention on the neural response to noise in the Attention Task. Electrode topographies are contralateralized to represent left side stimulation, and collapsed across noise frequencies (10 and 15 Hz). Stars indicate the cluster of electrodes that showed a significantly greater effect of attention during correct trials than incorrect trials (cluster-corrected *p* < .001).

### Invisible Signals Elicit Reliable Frequency Responses

A central goal of our study was to determine whether invisible (and visible) signals elicit reliable SSVEPs. To do this we calculated power at the signal frequencies (5 and 7.5 Hz, see Methods) and collapsed across frequencies and attention conditions. We then compared the electrode distributions to a zero power electrode distribution with a one-tailed Monte-Carlo cluster-based permutation test (between participant factor: electrode power, cluster *p* < .05, unit *p* < .05, 1000 permutations), separately for each level of awareness. As revealed in Figure 8, signal frequency power during presentation of a visible signal was significantly greater than zero across a broad posterior and mostly contralateral cluster of electrodes (cluster-corrected *p* = .002), confirming the presence of a neural response to visible signals. Crucially, signal frequency power during presentation of invisible signals was also significantly greater than zero across a cluster of posterior and mostly contralateral electrodes (cluster-corrected *p* = .002), confirming the presence of a neural response to invisible signals.

**Figure 8.**
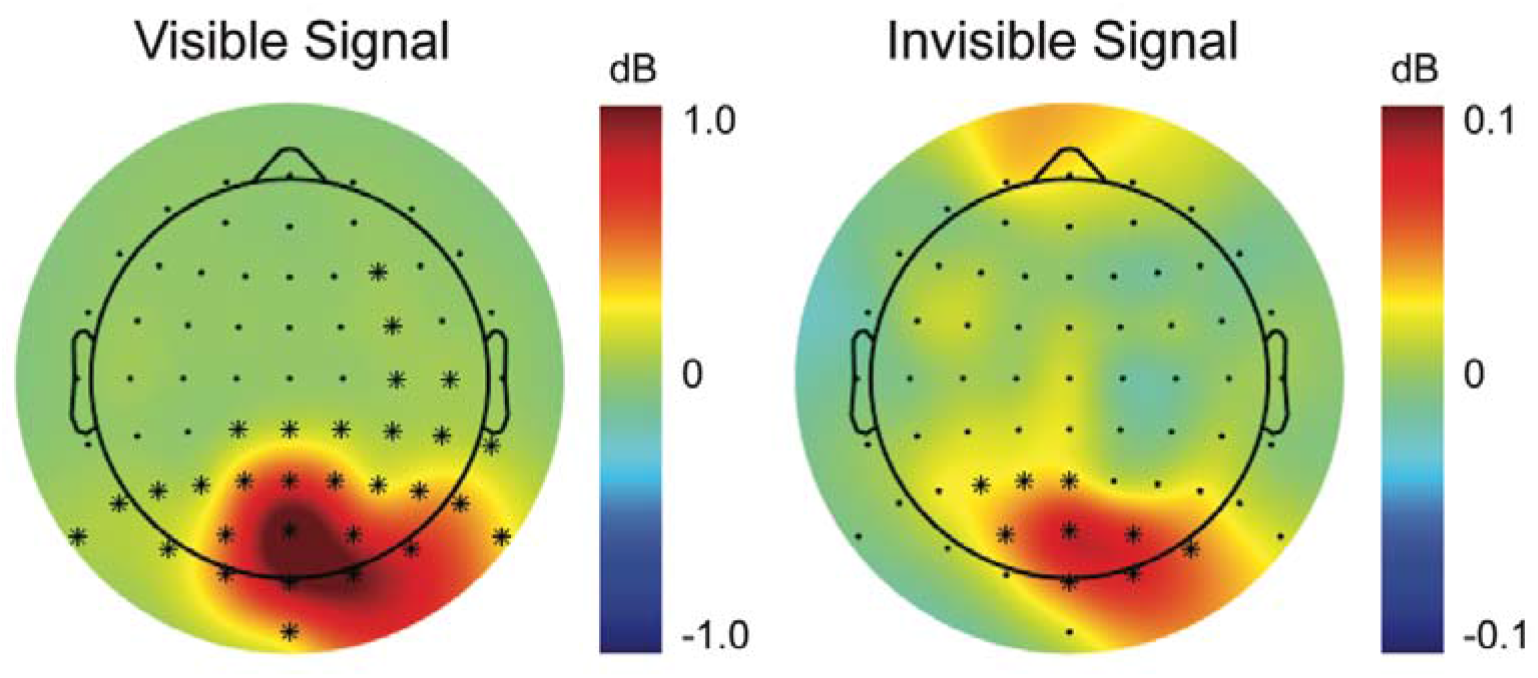
Neural response to visible and invisible signals in the Attention Task. Electrode topographies represent SSVEP power in response to visible signals (left) and invisible signals (right), contralateralized to represent left side stimulation, and collapsed across attention conditions and signal frequencies (5 and 7.5 Hz). Stars indicate clusters of electrodes with significant signal relative to a zero power topography map (cluster-corrected *p* < .05).

### Signal Frequency Responses Are Not Driven by Noise Stimuli

As a control, we checked whether the neural activity observed at signal frequencies might reflect a neural response to noise stimuli at half the frequency of stimulation. To do this we computed frequency power in Awareness Task intervals (see Methods) and collapsed across the cluster of electrodes that showed a significant response to invisible stimuli in the Attention Task (Pz, POz, Oz, PO3, PO4, contralateral PO7/8, contralateral O1/2, ipsilateral P1/2 and ipsilateral P3/4; see Figure 8), separately for intervals that contained signal and those that contained only noise (at each frequency of interest). As can be seen in Figure 9, Awareness Task intervals that contained signal (grey lines) elicited peaks in the frequency spectrum at signal frequencies (5 or 7.5 Hz), but intervals that contained only noise (black lines) produced no such activity. One-tailed *t*-tests demonstrated that signal frequency power was greater than zero during signal intervals (M = .08 dB, SD = .07 dB, *t* = 5.931, *p* < .001), but no greater than zero during noise-only intervals (M < .01 dB, SD = .01 dB, t = .965, *p* = .172). Crucially, Bayesian statistics supported the null hypothesis that noise stimuli produced no neural response at signal frequencies (uniform prior, lower bound = 0, upper bound = .08 dB, *B* = .10). This finding aligns with that of a previous study that used an analogous frequency tagging approach (Ales et al., 2012), which demonstrated no neural response at the signal frequency when participants were shown image sequences with noise images only. Together, these results confirm that the observed neural activity at signal frequencies in the Attention Task was driven by signal stimuli, and not a subharmonic response to noise stimuli.

**Figure 9.**
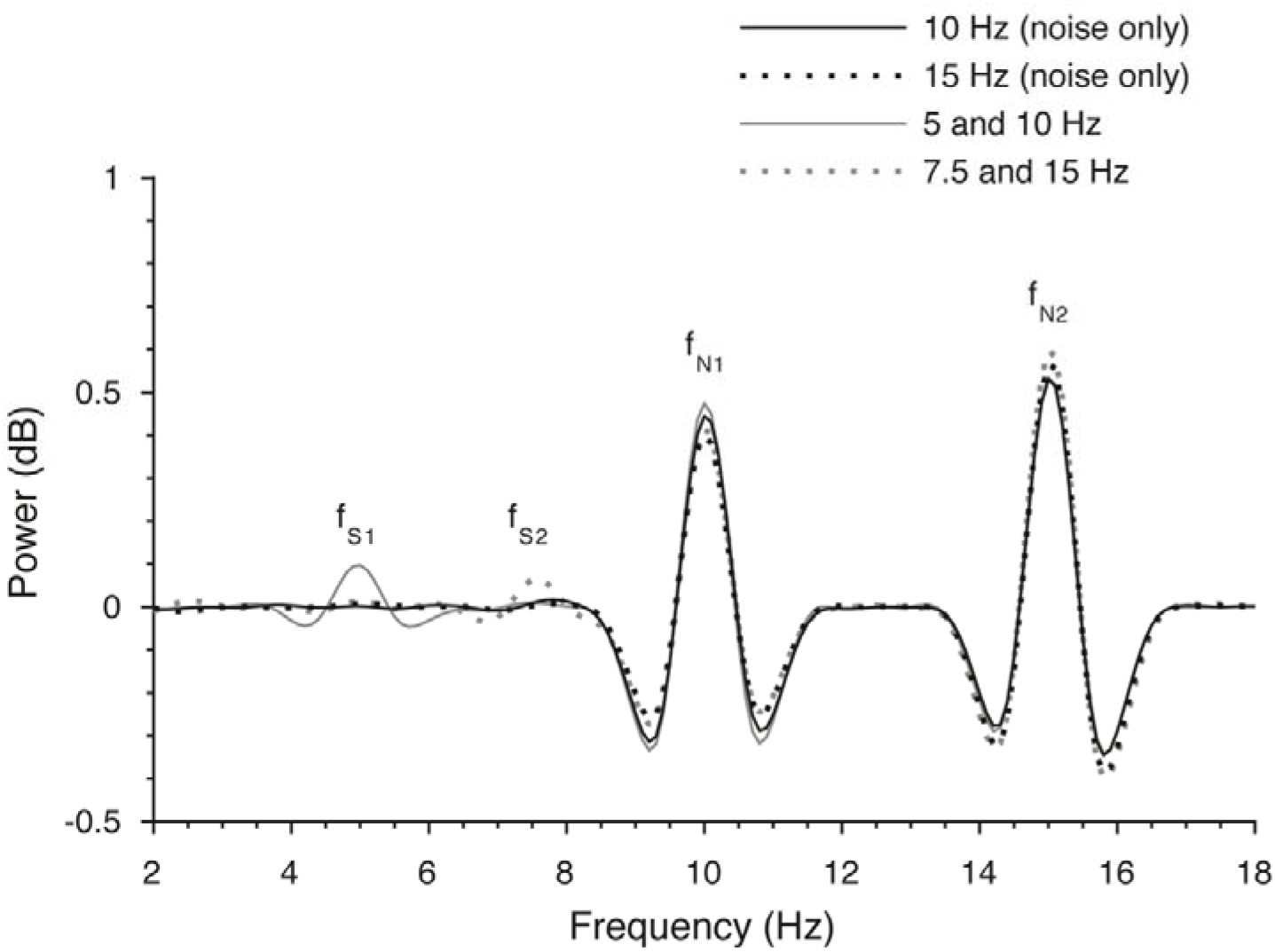
Phase-locked power in the Awareness Task, averaged across the cluster of electrodes that showed a significant response to invisible signal in the Attention Task (see Figure 8). Intervals that contained only noise at the frequency of interest are shown in black and intervals that contained both signal and noise are shown in grey. Note that noise-only intervals did not elicit peaks in the frequency spectrum at signal frequencies (5 and 7.5 Hz).

### Attention Enhances Neural Representations of Visible and Invisible Signals

Considering the weaker neural response to signals compared with high-contrast noise (Figure 5), we collapsed power across posterior and contralateral clusters of electrodes that showed a significant response to the signal (Figure 8), separately for each level of awareness and attention. As revealed in Figure 10, attention increased the neural response to both visible and invisible signals across these electrode clusters. A two-way analysis of variance tested the effects of signal coherence (two levels: *visible*, *invisible*) and spatial attention (two levels: *attended, ignored*) on neural responses to signal. Results of the ANOVA revealed a main effect of signal coherence (*F*_(1,22)_ = 47.699, *p* < .001 , *η_p_^2^* = .68), with greater neural responses to visible signals (M = .35 dB, SD = .25 dB) than to invisible signals (M = .05 dB, SD = .05 dB). Spatial attention also increased neural responses to stimuli (*F*_(1,22)_ = 7.693, *p* = .011, *η_p_^2^* = .26), with significantly greater signal frequency power in response to attended signals (M = .24 dB, SD = .16 dB) than ignored signals (M = .16 dB, SD = .16 dB). The interaction between signal awareness and spatial attention was also significant (*F*_(1,22)_ = 4.768, *p* = .040, *η_p_^2^* = .18). Follow-up paired-samples t-tests revealed that the interaction was driven by a greater effect of attention on visible signals (ΔM = .11 dB, ΔSD = .19 dB) than invisible signals (ΔM = .04 dB, ΔSD = .08 dB, *t*_(22)_ = 2.18, *p* = .040).

**Figure 10.**
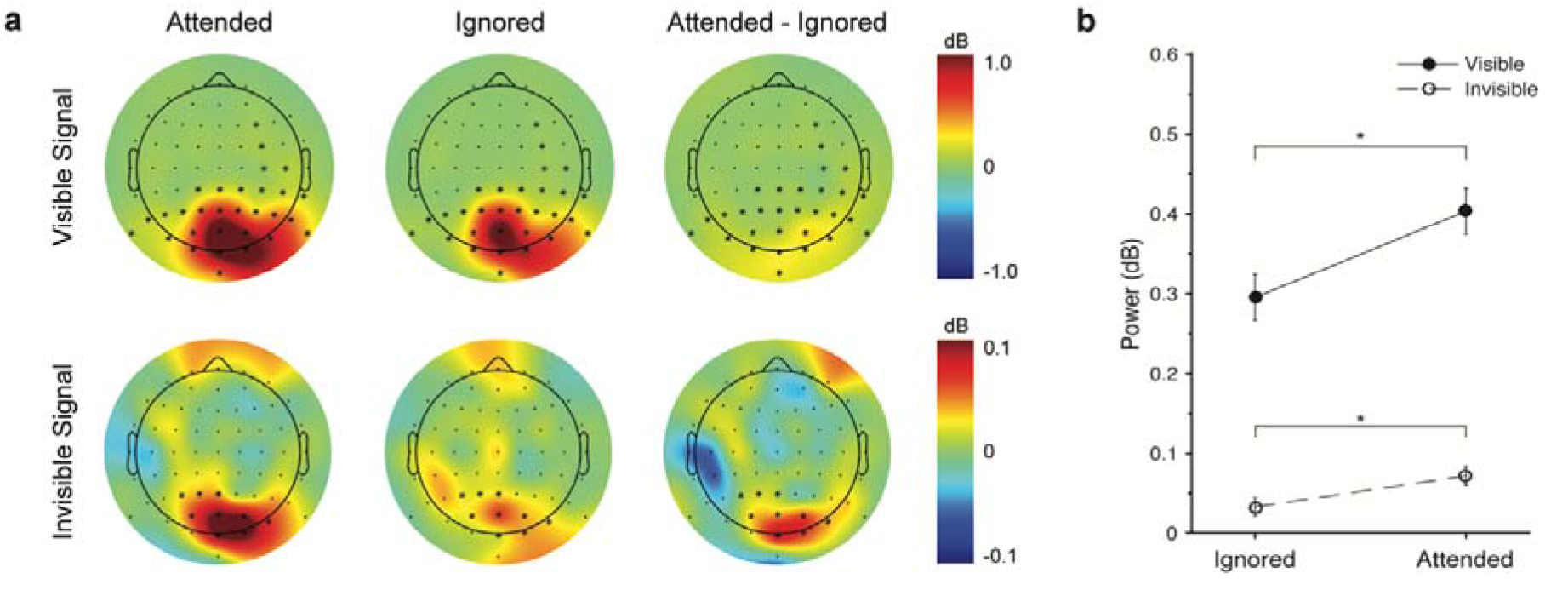
Effect of attention on neural responses to visible (top) and invisible (bottom) signals in the Attention Task. **(*a*)** Electrode power topographies for attended signals (left), ignored signals (middle), and the difference between attended and ignored signals (right). Topographies are contralateralized to represent left side stimulation, and collapsed across signal frequencies (5 and 7.5 Hz). Stars indicate the electrodes showing significant signal (Figure 8), across which power was collapsed to investigate the effect of attention. **(*b*)** Effect of attention within each level of awareness, collapsed across electrodes showing significant signal. Attention significantly increased the neural response to both visible and invisible signals (p < .05).

Since our critical research question related to whether attention can modulate neural responses to invisible stimuli, we also followed up the main effect of attention with t-tests of the simple main effect of spatial attention at each level of signal awareness (Figure 10b). Spatial attention enhanced neural responses to visible signals, with greater activity in response to attended (M = .40 dB, SD = .26 dB) than ignored visible stimuli (M = .30 dB, SD = .27 dB, *t*_(22)_ = 2.671, *p* = .014). This finding is in line with previous research showing attentional enhancement of SSVEPs to visible flickering stimuli (Vialatte, Maurice, Dauwels, & Cichocki, 2010). Crucially, spatial attention also modulated neural responses to invisible signals, with significantly greater activity in response to attended (M = .07 dB, SD = .07 dB) than ignored invisible stimuli (M = .03 dB, SD = .06 dB, *t*_(22)_ = 2.363, *p* = .027), indicating that attention can also enhance neural representations of invisible stimuli embedded in highly salient noise.

## Discussion

Previous research has suggested that covert spatial attention can modulate neural processing of invisible stimuli, supporting the notion that attention and awareness are dissociable neural mechanisms (Watanabe et al., 2011; Wyart et al., 2012; Wyart & Tallon-Baudry, 2008). Nevertheless, the intricacies of such a relationship remain poorly understood. In particular, no study to date has demonstrated that spatial attention can modulate *neural representations* of invisible stimuli, or assessed the nature of such modulation when those stimuli are in spatial competition with highly salient noise. To investigate these questions, we developed a novel attention task in which participants counted the number of brief contrast decreases in one of two image streams that contained both signals (visible or invisible) and noise. We isolated neural responses to noise in cued (attended) and non-cued (ignored) image streams, and observed enhanced activity across contralateral and posterior electrodes to cued noise throughout the trial epoch, confirming that participants voluntarily held their attention to one of the two lateralized image streams as instructed. The effect of attention on neural responses to noise was also greater across frontal and central electrodes with correct identification of contrast targets (Figure 7), suggesting that fluctuations in attention across trials directly affected target detection.

We employed a novel frequency tagging approach that allowed us to isolate neural representations of visible and invisible signals embedded in highly salient noise. To our knowledge, this is the first study to report SSVEP responses to objectively invisible stimuli embedded in noise. It could be argued that since we did not measure signal awareness during the Attention Task, participants might have been aware of the ‘invisible’ signal. Although we cannot rule this out, such a scenario is highly unlikely, considering that participants actively searched for signals in the Awareness Task, but looked instead for contrast decrements during the Attention Task. Thus, our results suggest that awareness of a masked stimulus is not a prerequisite for eliciting an SSVEP, as might be inferred from the step-like rise in SSVEP power that coincided with the onset of signal awareness in a previous study (Ales et al., 2012). Instead, our findings demonstrate that SSVEPs track intermediate levels of signal strength, even at levels too weak to provoke conscious perception.

Critically, our paradigm allowed us to measure the effects of spatial attention on neural representations of visible and invisible signals. We found that neural representations of visible signals were greater in the attended image stream than in the ignored stream, extending previous findings of effects of attention on neural representations of visible stimuli (Hillyard & Anllo-Vento, 1998; Martinez et al., 1999; Müller et al., 1998) to demonstrate that spatial attention also benefits partially degraded, yet still visible, signals in spatial competition with clearly visible and highly salient noise. Crucially, neural responses to invisible signals were also greater in the attended image stream than in the ignored stream, demonstrating that spatial attention enhances representations of invisible stimuli in direct spatial competition with highly salient and visible noise. Since spatial attention enhanced neural representations of signals without a corresponding increase in signal awareness, the present findings support the notion that spatial attention and awareness are dissociable neural mechanisms (M. A. Cohen, Cavanagh, Chun, & Nakayama, 2012; Dehaene, Changeux, Naccache, Sackur, & Sergent, 2006; Koch & Tsuchiya, 2012; Tallon-Baudry, 2012).

Although the present study is not the first to demonstrate effects of spatial attention in the absence of object awareness (Schurger et al., 2008; Watanabe et al., 2011; Wyart et al., 2012; Wyart & Tallon-Baudry, 2008), it makes several important advances on the existing literature. First, the present study investigated a distinct question to that of previous studies that aimed to assess the effects of both awareness and attention on neural processing, independent of signal strength (Schurger et al., 2008; Wyart and Tallon-Baudry, 2008; Wyart et al., 2012). In these previous studies, signals were presented at detection threshold, and participants’ subjective reports were used to categorise trials as visible or invisible. The authors found effects of attention on the neural processing of peri-threshold signals, even when participants reported being unaware of their presence. As such, these studies provide evidence that attention can modulate peri-threshold stimuli, but cannot speak to how the visual system treats very weak signals with insufficient bottom-up activation to enter awareness, irrespective of the cognitive state of the observer (so-called ‘subliminal’ stimuli, Dehaene et al., 2006). In the present study, we presented visible and invisible signals at different, pre-determined levels of coherence, and verified that invisible stimuli were objectively undetectable with a two-interval forced-choice signal detection task. Thus, we can be confident that the invisible stimuli in our experiment were not perceived due to a lack of bottom-up activation, rather than fluctuations in the cognitive state of the observer. Correspondingly, our findings demonstrate that neural processing of objectively subliminal stimuli can be modulated by spatial attention, as suggested by Dehaene and colleagues (2006), and that surpassing a hypothetical ‘threshold’ is not a necessary precursor for modulation by spatial attention.

Second, previous studies have not demonstrated that the observed neural activity, modulated by attention, was specifically related to the invisible stimuli in question. As such, previously observed effects of attention may instead reflect (a) baseline shifts in neuronal activity that occur even in the absence of external driving stimuli (as may be the case in Watanabe et al., 2011; see Driver and Frith, 2000) (b) enhanced neural representations of other, visible stimuli (e.g. the spatial cue in Wyart et al., 2012, as has been argued by Cohen et al., 2012), or (c) subcomponents of spatial attention that do not modulate neural representations *per se* (e.g. spatial re-orienting after a miscued stimulus in Schurger et al., 2008; Wyart and Tallon-Baudry, 2008; Wyart et al., 2012). In demonstrating that spatial attention modulates specific neural correlates of invisible stimuli, without a corresponding increase in awareness, the present study provides the first clear evidence that spatial attention and awareness dissociate at the level of neuronal representations.

A third, and arguably most important, advance of the current study is that we have shown that spatial attention can enhance neural representations of invisible stimuli that are in direct spatial competition with highly salient, visible stimuli. Previous studies presented invisible signals alone (Schurger et al., 2008; Wyart & Tallon-Baudry, 2008), or at different times or locations (Watanabe et al., 2011; Wyart et al., 2012) to the salient masks used to titrate signal awareness. Since neural competition is maximal at the level of the receptive field (Beck & Kastner, 2009; Reynolds, Chelazzi, & Desimone, 1999), neural representations of invisible signals in these studies were likely under conditions of minimal competition. In contrast, we maximised competition between signal and noise by presenting them concurrently and at the same location. Our findings reveal concurrent neural representations of both visible and invisible stimuli at the same location, demonstrating that spatial competition with highly salient stimuli is not sufficient to suppress weak neural representations of invisible stimuli. Moreover, the present study demonstrates that weak neural representations of invisible stimuli in competition with salient stimuli can nevertheless be biased according to the top-down goals of the observer – in this case, holding covert attention preferentially to the left or right visual field. Given that signal features were irrelevant to the contrast detection task, this finding suggests that all stimuli at attended locations are prioritised relative to those at unattended locations, irrespective of their task-relevance, their capacity to enter awareness, or their proximity to more salient stimuli.

The present findings demonstrate that spatial attention can operate independent of mechanisms of awareness, at the level of neural representations. More broadly, the present findings place spatial attention within a growing body of literature that suggests various forms of attention (e.g., temporal, feature-based, and involuntary spatial attention) can operate in the absence of stimulus awareness (for a review, see Koch & Tsuchiya, 2007). Together, these findings argue against the idea that attention and awareness are identical (Prinz, 2012) and instead support theories that cast attention and awareness as dissociable mechanisms (M. A. Cohen et al., 2012; Dehaene et al., 2006; Koch & Tsuchiya, 2012; Tallon-Baudry, 2012). Nevertheless, the exact nature of this relationship remains to be fully characterized, in particular whether the different forms of attention interact with awareness according to the same underlying principles, and how such top-down biases interact with bottom-up processes related to salience and neural competition between representations. To this end, we anticipate that the present paradigm could be adapted to study how other non-spatial forms of attention (e.g., feature-based attention) modulate neural representations of multiple competing stimuli at different levels of awareness.

## Supplementary Material

**Movie 1.** Example trial of the Awareness Task. At the beginning of the trial, static noise images appear on either side of fixation, and central arrows indicate the image stream to be attended (in this example, the left stream). After 0.5 *s* the image streams flicker for the first 2.4 *s* interval, are static for another 0.5 *s*, and then flicker again for the second 2.4 *s* interval. On the cued (left) side, one of the two flickering intervals contains signal embedded in every second image (in this example, the second interval), the coherence of which increases linearly during the first 0.4 *s* of the interval. Signal is also present in one of the two intervals on the non-cued (right) side (in this example, the first interval).

**Movie 2.** Example trial of the Attention Task. At the beginning of the trial, static noise images appear on either side of fixation, and central arrows indicate the image stream to be attended (in this example, the left stream). After 0.5 *s* the image streams flicker for 10.4 *s*. At the end of the trial participants report how many times the cued (left) image stream decreased in contrast (in this example, twice). The non-cued image stream also contains up to two contrast decrements (two in this example). Both image streams contain signal embedded in every second image, the coherence of which increases linearly during the first 0.4 *s* of flicker to a level that is either visible or invisible to the participant (in this example, the left image stream contains visible signal and the right image stream contains invisible signal).

